# Hippocampal circuit abnormalities in MeCP2+/− mouse model of Rett syndrome

**DOI:** 10.1101/034835

**Authors:** Julia Chartove, Wenlin Liao, Aniqa Hassan, Mary McMullen, Rachel White, Sangwon Kim, Gregory C. Carlson

## Abstract

Rett syndrome (RTT) has a complex developmental course over childhood and adolescence. Patients with RTT often have a pre-symptomatic period with no or little outward signs of the disorder, followed by developmental arrest and regression. Following regression, the individual’s condition is not static, as they often progress into defined stages with unique neurological symptoms. Similarly, the progression of RTT-like symptoms in female mice heterozygous for a null-mutation has a prodromal and symptomatic period. Change in functional local circuit connectivity was studied using hippocampal slices, assaying Schaffer evoked activity in area CA1 using fast voltage sensitive dye imaging. With this technique the local functional interactions between the excitatory and inhibitory components of the circuit can be characterized. The prodromal period was associated with a shift in extent of excitation into the stratum oriens of the hippocampus and reduced sensitivity to changes in divalent cation concentration. These data suggest that hyperexcitability of the hippocampus at the circuit level may contribute to the prodromal reduction in cognitive performance and the onset of developmental regression.

## Introduction

Rett Syndrome is a rare developmental disorder affecting an estimated 1 in 10,000 girls and women worldwide. It is characterized by the development of a constellation of motor and behavioral symptoms, such as difficulty or inability to walk, ataxia, lack of social reciprocity or use of nonverbal social behavior, lack or loss of speech, and deficits in sensory integration. In addition, Rett Syndrome is often (in up to 80% of cases) comorbid with epilepsy. (Tetzchner, 1996; Charour, 2007) Rett Syndrome is typically caused by a de novo mutation of the methyl CpG binding protein 2 (MECP2) gene on the X chromosome. The protein encoded by MECP2 is necessary for inactivating certain methylated genes, and is expressed constitutively but is found in particularly high concentrations in the brain. (Amir, 1999) Because MECP2 is located on the X-chromosome, Rett syndrome is a sex-linked disorder; male humans with no functional copies of MECP2 tend to be stillborn, so Rett syndrome is typically not diagnosed in these individuals. (Chahrour, 2007) In girls, however, symptoms of Rett syndrome do not appear until between 6 and 18 months of age, at which point there begins a pattern of developmental regression similar in time scale to that seen in autism. (Neul, 2004)

MECP2 exists on the X chromosome of mice as well as humans, meaning that female mice heterozygous for MECP2 are an appropriate animal model for Rett Syndrome. (Ezeonwuka, 2014; Ricceri, 2008) Indeed, these mice have been shown to replicate behavioral and physiological phenotypes seen in humans with Rett syndrome. MECP2 knockout and heterozygous mice have a similar pattern of developmental regression to the one seen in human Rett patients, after which they display symptoms of anxiety and behavioral stereotypies, as well as deficits in learning and memory. (Guy, 2001; Moretti, 2006; Pelka, 2006; Stearns, 2007) These symptoms are associated with decreased volume of the hippocampus, amygdala, and striatum, as well as impairment of synaptic plasticity and long-term potentiation in the hippocampus. (Pelka, 2006; Stearns, 2007; Asaka, 2006; Wither, 2013) Unlike humans, male MECP2 knockout mice do survive for several months, so they are also of interest as Rett syndrome models with a more severe phenotype. (Ricceri, 2008) Several studies in our lab and others have found that Rett mouse models show disruptions of typical oscillatory activity; of particular interest is the increased power of gamma frequency oscillations, especially in the hippocampus. (Liao, 2012; Goffin, 2012; D'Cruz, 2009) These abnormalities may underlie the deficits in learning, memory, and sensory processing seen both in Rett mouse models and Rett patients, as well as playing a role in the frequent comorbidity of epilepsy. (Uhlhaas, 2011; McLeod, 2013) These findings are of interest not only for their relationship with mechanisms of cognitive deficits in Rett syndrome, but also their relevance to autism and schizophrenia, in which disruptions in gamma oscillations have been implicated. (Uhlhaas, 2010; Wang, 2010)

Though before 4 months of age, RTT model mice do not show obvious symptoms, several studies have identified subtle behavioral abnormalities such as anxiety-like behavior, motor deficits, and sleep apnea in prodromal mice. (De Filippis, 2010; Ricceri, 2008; Ren, 2012) Additionally, prodromal RTT mice have been shown to have neurobiological abnormalities in their dendritic spine density, GABAergic synapses, protein synthesis capacity, and neuronal excitability. (Chapleau, 2012; Medrihan, 2008; Ricciardi, 2011; Gantz, 2011) However, aside from Isoda et al’s 2010 study that showed deficits in serotonergic innervation to the hippocampus of pre-symptomatic RTT mice, little work has been done identifying the characteristics of the Rett prodrome on a circuit level. Our study seeks to elaborate upon these findings by examining MECP2+/− female mice at 1 to 4 months of age, before Rett-like symptoms have begun to develop. By doing so, we hope to characterize the origins of the developmental deficits that lead to later pathology. We chose to utilize voltage sensitive dye imaging (VSDi) of slices taken from the hippocampus of 1 to 2 month old prodromal MECP2+/− female mice, 3 to 4 month old prodromal MECP2+/− female mice, and 1 to 2 month old symptomatic MECP2-null male mice, as well as age and sex matched wild type controls. When the Schaffer collaterals are directly stimulated, CA1 in hippocampal slices exhibits a pattern of excitation followed by rapid inhibition that is believed to be an underlying mechanism of gamma oscillatory activity. (Buzsaki, 2012; Whittington, 2010) Utilizing VSDi, we hope to characterize the intensity, spatial extent, and time course of both components of this response in each age and sex group and determine how they are affected by the MECP2 genotype.

## Methods

### Experimental setup

Wild-type littermates (WT) or Rett model (RTT) mice were decapitated, their brains removed and blocked in ice cold artificial cerebrospinal fluid (ACSF) with NaCl replaced with an equal osmolar concentration of sucrose. For each slice, five trials were conducted in typical ACSF solution and five trials were conducted in ACSF containing a high concentration of divalent cations (hi-di solution). These divalent cations raise the voltage of the extracellular medium and therefore prevent depolarization that results from opening of neurotransmitter-gated ion channels, but do not prevent depolarization generated by direct electrical stimulation; therefore, hi-di solution prevents only polysynaptic activation. (Liao, 2002; Sivaramakrishnan, 2013)

Hippocampal slices (350 μm) were cut at 12° off horizontal. Slices were allowed to recover in a static interface chamber at 35 °C for 30 min and then stored at room temperature. For voltage sensitive dye recordings, slices were stained with 0.125 mg/ml JPW 3031 in ACSF, and imaged in an oxygenated interface chamber using a fast 80X80 CCD camera recording at 2 kHz. (NeuroCCD, Redshirt Imaging, CT) For further details, see Nature Protocols. (Carlson, 2008) Stimulation was via a bipolar electrode placed in the Schaffer collaterals. Each slice underwent 10 stimulations; 5 in regular ACSF at 20 microamperes (μA), 12 μA, 8 μA, 4 μA, and 20 μA again, then 5 stimulations of the same intensities in hi-di solution.

### Image processing

All analysis was performed in IGOR (Wavemetrics, OR) on video data generated by averaging 12 trials of equal stimulation intensity. The overall fluorescence intensity of each pixel of the greyscale image was first normalized to that of the period immediately prior to stimulation. To compensate for photobleaching of the dye, fitted single exponentials were subtracted from these normalized fluorescence traces, which were plotted as change in fluorescence over total fluorescence (ΔF/F). A spatially segregated region in which no electrical activity was observed was selected for each slice as a null baseline, and the average intensity in this region was subtracted from the entire image to reduce background noise. To smooth the images while maintaining their statistical properties, median filtering was applied, both 2-dimensional 3×3 pixel filtering of each frame and 3-dimensional 3×3×3 pixel filtering of 3 adjacent movie frames. Images were then colorized according to fluorescence intensity on a “rainbow” spectrum from red (representing −0.0005 ΔF/F) to blue (representing 0.0003 ΔF/F). Red and blue were chosen in order to clearly visually correspond to excitatory and inhibitory neural responses, since the dye increases in fluorescence in response to a negative change in charge and decreases in fluorescence in response to a positive change in charge.

Because each video consisted of 1024 80 by 80 pixel frames, and each individual slice was tested at several stimulus intensities in both regular ACSF and high divalent cation solution, the volume of data generated for this study necessitated that analysis be automated. The analysis detailed above was completed for each slice once, and two regions of interest corresponding to the stratum radiatum and stratum oriens were outlined. We designed an analysis package, NP2Beta, to then repeat these analyses for all trials performed on the same slice and generate various graphical and numerical summaries of the analysis results. Along with the colorized and filtered video, these summaries included two types of traces that plotted activity over time: graphs of average change in fluorescence in an area (stratum radiatum or stratum oriens) and graphs of the percentage of an area over or under an intensity threshold (0.0003 ΔF/F or −0.0005 ΔF/F, respectively). The latter measure is referred to in this paper as “percent activation”. The times of peak excitation and peak inhibition within each area were identified by locating the minimum of the fluorescence intensity trace over the 350–380 millisecond timeframe and the maximum over the 350–480 millisecond timeframe, respectively (since stimulation occurred 350 milliseconds into the recording). These time points and the corresponding fluorescence intensity and percentage over/under threshold values were tabulated for each set of twelve trials for use in statistical analysis.

## Results

To test for differences in hippocampal excitability, we applied electrical stimulation to the Schaffer collaterals of hippocampal slices of 1 to 2 month old MECP2+/− female mice, 3 to 4 month old MECP2+/− female mice, and 1 to 2 month old MECP2-null male mice, and recorded gross electrical response via voltage sensitive dye imaging. Within CA1, the stratum radiatum and stratum oriens were analyzed separately in terms of both affected area (percent activation) and intensity of the excitatory and inhibitory phases of the response, as measured by change in dye fluorescence. In addition, during some trials ACSF containing a high concentration of divalent cations was applied in order to diminish capacity for polysynaptic response and isolate only those cells responding directly to the input from the electrode.

The percentage of the stratum oriens displaying electrical excitation generating greater than −0.0005 ΔF/F was significantly greater in 3–4 month old RTT females and 1–2 month old RTT males than in age and sex matched wild type controls. (Figure 1) In 3–4 month old wild type females, the mean percent activation in stratum oriens at the time of peak excitation was 18.24%, while in RTT females, it was 31.40%. (mixed-model nested ANOVA, F(1, 10) = 11.52, p = 0.0012*) In 1–2 month old males, this difference is even more pronounced, with wild type males having a mean of only 10.43% activation in oriens and RTT males showing 35.13% activation. (F(1,7) = 9.61, p = 0.0044*) This difference is present in the younger females but much smaller, with 1–2 month old wild type females and RTT females having no significant difference in percent activation in oriens; on average, they showed 23.12% activation and 17.33% activation, respectively, indicating that the magnitude of the observed differences in oriens depend on both age and sex. (F(1, 10) = 2.94, p = 0.093) No significant differences in extent of activation in the stratum radiatum were observed in any age or sex group. Furthermore, no significant differences in extent of inhibition in any area were observed, indicating that these differences in excitation are not related to a deficit in the spatial distribution of the inhibitory component of the response in RTT mice. (Table 1)

**Figure 1.**
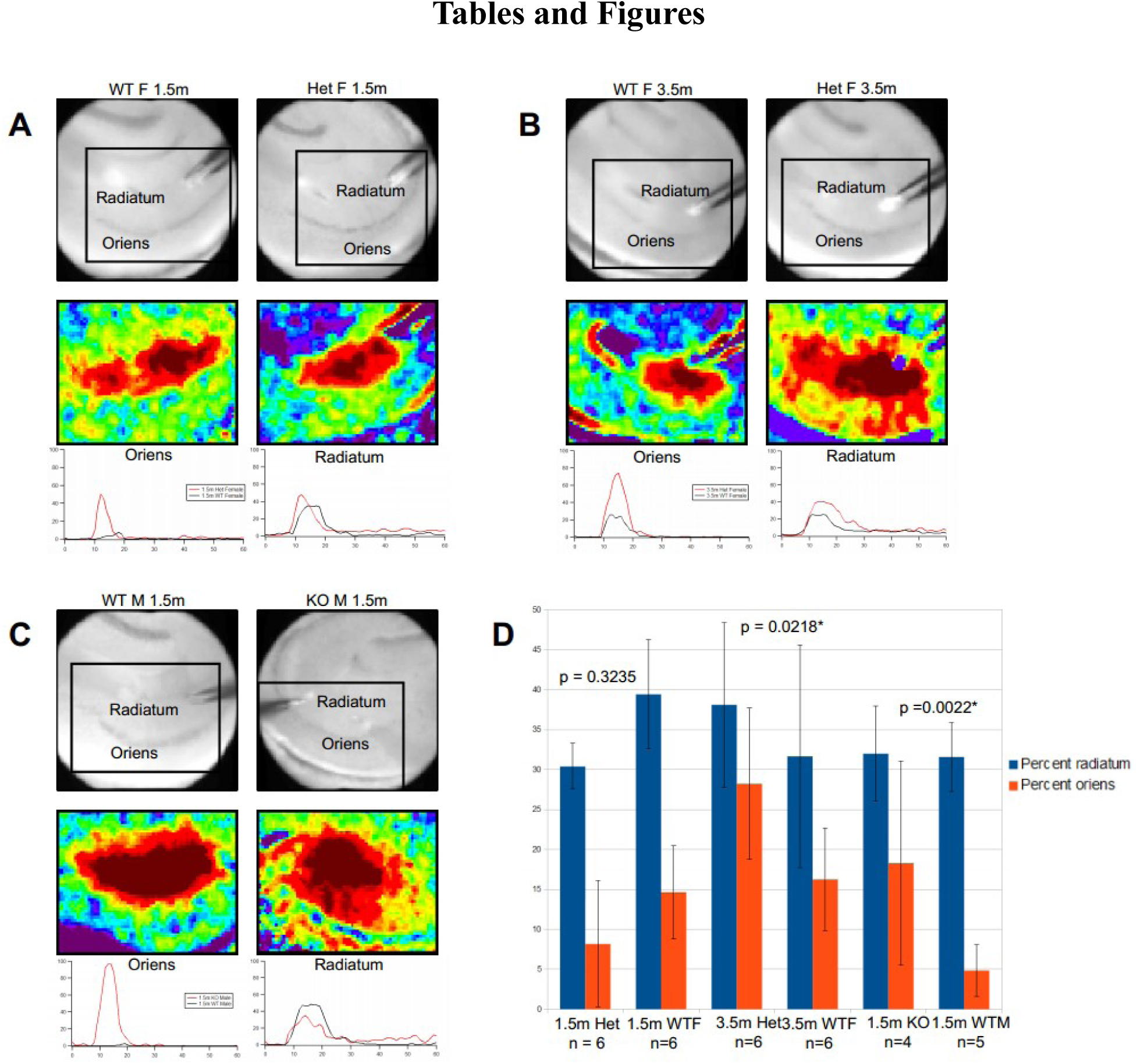
**A**. Top: Grayscale images of slices taken from female WT and RTT mice at age 1.5 months. Middle: Colorized VSDi images corresponding to activation in regions delineated by boxes. Bottom: Graphs of percentage of radiatum and oriens under −0.0006 ΔF/F in each slice over time. **B**. Slices taken from female WT and RTT mice at age 3.5 months. **C**. Slices taken from male WT and RTT mice at age 1.5 months. Note differences between extent of activation in oriens in WT (black) and RTT (red) individuals versus differences between extent of activation in radiatum. **D**. Median percent activation within radiatum and oriens of each group. Error bars represent standard error. p values refer to Wilcoxon rank-sum tests of oriens excitation. Excitation in oriens is significantly greater in 3.5m RTT females and 1.5m RTT males than in age and sex matched wild types.

**Table 1:** Percent activation by sex, age, solution, genotype, and brain region.

| Sex | Age | Solution | Genotype | Oriens<br>excitation | Oriens<br>inhibition | Radiatum<br>excitation | Radiatum<br>inhibition |
| --- | --- | --- | --- | --- | --- | --- | --- |
| Female | 1.5m | ACSF | WT | 23.128 | 47.836 | 36.462 | 42.604 |
|  |  |  | RTT | 17.332 | 39.706 | 31.453 | 35.588 |
|  |  | Hi-di | WT | 18.536 | 55.241 | 32.448 | 42.344 |
|  |  |  | RTT | 17.706 | 38.473 | 37.993 | 39.162 |
|  | 3.5m | ACSF | WT | 18.238 | 43.567 | 44.855 | 39.379 |
|  |  |  | RTT | 31.398 | 38.206 | 42.446 | 36.829 |
|  |  | Hi-di | WT | 10.903 | 40.846 | 29.863 | 36.172 |
|  |  |  | RTT | 26.039 | 37.448 | 33.244 | 35.557 |
| Male | 1.5m | ACSF | WT | 10.428 | 40.765 | 31.384 | 36.403 |
|  |  |  | RTT | 35.133 | 38.580 | 36.368 | 27.996 |
|  |  | Hi-di | WT | 10.061 | 41.306 | 28.246 | 32.431 |
|  |  |  | RTT | 47.362 | 44.549 | 42.901 | 29.824 |

After bath application of ACSF containing higher concentrations of divalent cations, 10 of 12 WT individuals showed a decrease in peak intensity of excitation, as hypothesized, with the other two individuals showing marginal increases. On average, the decrease in intensity of excitation in stratum radiatum when comparing the first stimulation in regular ACSF with the first stimulation in hi-di solution was 8.628×10^−5^ ΔF/F(matched pairs t(11) = 2.99, p = 0.012*). In contrast there was no group level change in excitability in the RTT mice; on average, the effect of hi-di solution was to produce a marginal increase of 2.273×10^−5^ ΔF/F in peak intensity. (t(10) = −0.0006, p = 0.999). These patterns were observed within all age and sex groups. (Figure 2) The stratum oriens showed the same trend, with peak excitation in WT individuals decreasing by 1.658×10^−5^ ΔF/F but peak excitation in RTT individuals increasing by 6.209×10^−5^ ΔF/F, though these trends were not significant (p = 0.227 and p = 0.417, respectively). High divalent cation solution did not have any significant impact on the inhibitory component of the response, indicating that most of the inhibition generated by stimulation was via direct activation of inhibitory neurons. (Table 2)

**Figure 2.**
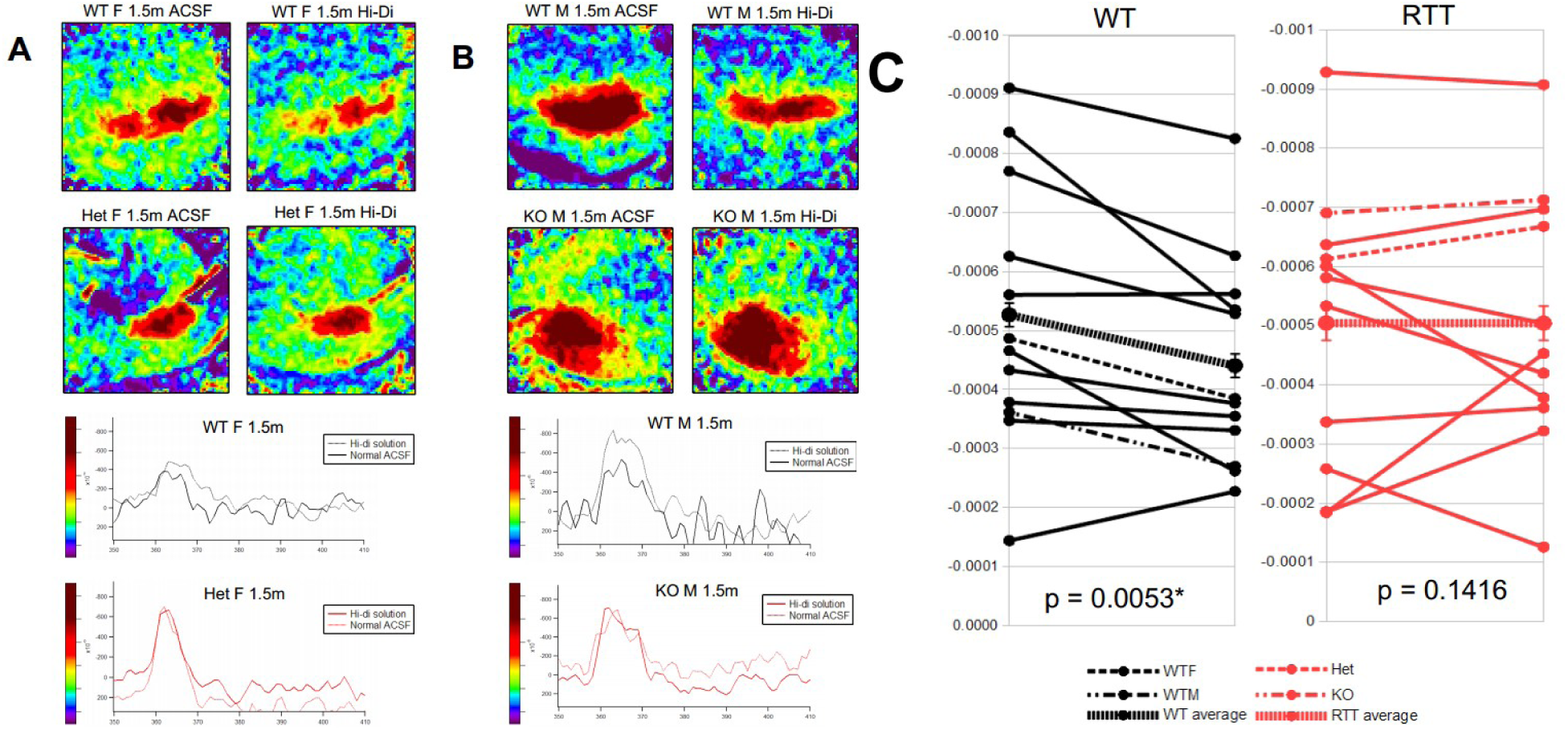
**A**. Top: Colorized VSDi images of slices taken from WT (top row) and RTT (second row) females at age 1.5 months. Within each pair, images on the left are from trials using normal ACSF; images on the right are from trials using high divalent cation solution. Bottom: Graphs of average intensity of activation over time within radiatum of each slice. **B**. Slices taken from WT and RTT males. Note that in both groups, WT individuals (black) show differences in peak intensity between normal ACSF (dotted lines) and hi-di solution (solid lines); RTT individuals (red) do not. **C**. Matched pairs of peak intensities between normal ACSF and hi-di solution trials for each slice. 10 of 12 WT individuals show a decrease in intensity in hi-di solution, as opposed to only 5 of 11 RTT individuals. *Dotted lines represent the examples from A and B*.

**Table 2:** Peak intensity (ΔF/F) by sex, age, solution, genotype, and brain region.

| Sex | Age | Solution | Genotype | Oriens<br>excitation | Oriens<br>inhibition | Radiatum<br>excitation | Radiatum<br>inhibition |
| --- | --- | --- | --- | --- | --- | --- | --- |
| Female | 1.5m | ACSF | WT | $3.664 \times 10^{-4}$ | $2.422 \times 10^{-4}$ | $6.288 \times 10^{-4}$ | $2.067 \times 10^{-4}$ |
| | | | RTT | $3.299 \times 10^{-4}$ | $3.109 \times 10^{-4}$ | $5.980 \times 10^{-4}$ | $2.499 \times 10^{-4}$ |
| | | Hi-di | WT | $2.796 \times 10^{-4}$ | $2.542 \times 10^{-4}$ | $5.420 \times 10^{-4}$ | $2.002 \times 10^{-4}$ |
| | | | RTT | $3.175 \times 10^{-4}$ | $1.785 \times 10^{-4}$ | $5.165 \times 10^{-4}$ | $2.051 \times 10^{-4}$ |
| | 3.5m | ACSF | WT | $3.150 \times 10^{-4}$ | $2.018 \times 10^{-4}$ | $3.553 \times 10^{-4}$ | $1.545 \times 10^{-4}$ |
| | | | RTT | $3.603 \times 10^{-4}$ | $1.816 \times 10^{-4}$ | $5.386 \times 10^{-4}$ | $1.959 \times 10^{-4}$ |
| | | Hi-di | WT | $3.154 \times 10^{-4}$ | $1.809 \times 10^{-4}$ | $3.052 \times 10^{-4}$ | $1.368 \times 10^{-4}$ |
| | | | RTT | $4.431 \times 10^{-4}$ | $2.111 \times 10^{-4}$ | $5.020 \times 10^{-4}$ | $1.892 \times 10^{-4}$ |
| Male | 1.5m | ACSF | WT | $2.446 \times 10^{-4}$ | $2.187 \times 10^{-4}$ | $5.960 \times 10^{-4}$ | $1.465 \times 10^{-4}$ |
| | | | RTT | $8.610 \times 10^{-4}$ | $2.336 \times 10^{-4}$ | $3.533 \times 10^{-4}$ | $1.767 \times 10^{-4}$ |
| | | Hi-di | WT | $2.777 \times 10^{-4}$ | $1.628 \times 10^{-4}$ | $4.740 \times 10^{-4}$ | $1.850 \times 10^{-4}$ |
| | | | RTT | $1.197 \times 10^{-4}$ | $5.518 \times 10^{-4}$ | $4.959 \times 10^{-4}$ | $1.554 \times 10^{-4}$ |

## Discussion

Our findings indicate that significant abnormalities exist in the hippocampal circuitry of individuals with Rett syndrome even before any major behavioral differences are apparent. The early onset of differences in excitability and timing as well as the fact that these differences appear to worsen between the age of 1.5 months and 3.5 months suggests a potential causal role for these abnormalities in the developmental trajectory seen in Rett syndrome. Our results are in accord with prior research that has found hyperexcitability in Rett syndrome, both in cortex and in hippocampus. (Yamanouchi, 1993; Zhang, 2008) By isolating hippocampal differences to stratum oriens this work suggests that the direct entorhinal and thalamic inputs to that area may be primarily affected in this model.

There are a number of circuit level or intrinsic cellular differences that could drive this increase in excitability. Our results indicate that the hyperexcitability does not arise from intrinsic properties of MECP2-deficient neurons, but rather from hypoconnectivity that suggests a lack of inhibitory input and an imbalance between excitation and inhibition. (Dani, 2005; Zhang, 2008; Shepherd, 2011). High divalent cation solutions suppress polysynaptic activity to reduce evoked activity; in the MeCP2+/+ mice, normal polysynaptic connections that engage local circuitry are functional, resulting in suppression of activity in high-divalent cation-containing ACSF. This suppression is absent in the MeCP2+/− mice, indicating that at this stage of development these local circuit connections that drive normal hippocampal network activity are not functioning. This finding goes beyond a simple disruption of inhibitory/excitatory balance and supports an absence of normal local circuit functional connectivity during the prodromal period.

Hyperexcitability has been hypothesized to be the mechanism by which Rett syndrome impairs synaptic plasticity in hippocampus and therefore interferes with learning and memory; increased excitability reduces the dynamic range of synapses, therefore generating a ceiling effect wherein synapses are constantly potentiated to their maximum. (Weng, 2011; Calfa, 2011) However, the prodromal mice from which these slices were taken had yet to show significant memory impairments. Perhaps, though hyperexcitability was present in these mice, their synapses had not yet been potentiated to the degree that a ceiling effect could take place. This hypothesis could help explain why the developmental trajectory of regression in Rett syndrome only occurs after several months in both mice and humans, since the underlying cellular physiology is similar between species and therefore the degenerative loss of plasticity could occur at a similar rate.

The fact that spatial abnormalities in hyperexcitability could be localized specifically to the stratum oriens showcases the power of VSDi to pinpoint spatial differences with resolution and detail that are difficult to achieve using other imaging methods. The interneurons of the stratum oriens have been hypothesized to play an essential role in sensory signal selection and in generation of both gamma and theta oscillations, potentially helping to explain the sensorimotor deficits seen in Rett syndrome. (Gloveli, 2005; Kitchigina, 2010; Maccaferri, 2005) In addition, higher theta amplitudes have been shown to modulate gamma power, so disruptions in theta may help explain the abnormally high gamma power characteristic of Rett syndrome. (Canolty, 2006) Therefore, mechanisms of theta oscillations in the hippocampus of Rett model mice are of interest as an avenue of further research. Because we found the abnormal activity in slices from prodromal Rett mice to be localized to a specific spatial region, our findings suggest that through high-resolution imaging, it may be possible to detect Rett syndrome and potentially other disorders characterized by developmental regression in humans so that treatment may begin before the onset of symptoms. These applications would require further research into whether abnormalities detected using optical imaging could be used for diagnosis in awake, behaving animals, and whether non-invasive techniques could be used in place of direct optical methods for the purpose of diagnosis.

An important limitation of voltage sensitive dye that must be acknowledged is that the fluorescence signal produced by the dye represents a spatial average of all the cells within that region of the slice. Not every adjacent neuron has the same membrane potential at the same time, but we can only record one value for each pixel’s worth of area. Therefore, when we characterize activity as “excitatory” or “inhibitory” based on its overall fluorescence, there may well be activity of the opposite valence in the same spatial location that cannot be detected. This consideration becomes important when speculating on the mechanism of the patterns of hyperexcitability that we have observed in this study. It could be the case that in RTT mice, more excitatory neurons within the oriens are responding to the same level of stimulation; however, it could also be that within wild type mice, the signal from excitatory activity within the oriens is overwhelmed by inhibitory activity that is not present within RTT mice. Because the signal generated by the inhibitory response is weaker than the excitatory signal, the trends within the data we collected on inhibition may not have reached significance; there is in fact a general trend of weaker inhibition within the RTT mice as compared to the WT mice, as seen in Tables 1 and 2, so this possibility cannot be ruled out.

Another difficulty posed by utilizing voltage sensitive dye is analyzing the amount of data produced. Much of this difficulty can be overcome using computational methods to automate filtering and processing of data and to generate summary variables, as described above. However, this still leaves the task of deciding which summary variables to generate, and which trends within these variables to identify for further analysis. In addition to spatial extent and intensity of both excitation and inhibition within radiatum and oriens, data were collected on timing of peak spatial extent and of intensity of activation in both valences and both areas, as well as data on differential responses to different levels of stimulation. The findings reported in this paper were selected because they are characterized by robustness in terms of being present in multiple age and sex groups and existing as trends across multiple measures and stimulus intensities.

However, the choice of which findings to report ultimately did have a degree of subjectivity, and several intriguing patterns existed within the data but ultimately did not hold up to statistical scrutiny. In particular, we had an interest in how the timing of the inhibitory response differed between RTT and WT mice and whether this could impact the observed disruption in gamma oscillations. We did in fact find differences in inhibitory timing between genotypes, but the direction of these differences varied between age groups, sex groups, and hippocampal region. It could very well be that there is a complex pattern in which inhibitory timing is disrupted in certain populations of RTT mice, but it is more likely the case that the observed trends were due to chance variations as a result of noise overwhelming the already faint inhibitory response signal. Therefore, we chose not to report these trends in the results, but if we can obtain a cleaner signal they may be of interest for future research. Another set of data we attempted to collect was the decay time constant between the excitatory response and the inhibitory response, since we believed this also might be informative in terms of disruptions of gamma oscillatory timing. However, this measure was discarded due to difficulty in automating its collection; again, this issue could potentially be solved if the signal-to-noise ratio were improved, since a clearer signal would make generating an exponential fit easier. Potential techniques for improving signal to noise ratio in VSDi are described by Reynaud et al. (2011)

An interesting avenue for further research would be to perform the same manipulations on hippocampal slices from older female mice who are already displaying full RTT symptoms and to determine whether they display the same patterns of hyperexcitability. Several such experiments were attempted by our lab; however, they showed a great deal of variability which might be due to greater variability of a complex disease progression and rendered most of the data unusable. This window into the prodromal period does provide insight into a likely critical disruption of local circuit development.

## Acknowledgments

This research was supported by NIH/NICHD grant 1R01HD062577 and 1-P50-MH-096891–01. We would like to thank Kathleen Siwicki for her guidance in project management.

## Works Cited

Amir, Ruthie E., et al. “Rett syndrome is caused by mutations in X-linked MECP2, encoding methyl-CpG-binding protein 2.” Nature genetics 23.2 (1999): 185–188.

Asaka, Yukiko, et al. “Hippocampal synaptic plasticity is impaired in the Mecp2-null mouse model of Rett syndrome.” Neurobiology of disease 21.1 (2006): 217–227.

Buzsáki, György, and Xiao-Jing Wang. “Mechanisms of gamma oscillations.” Annual review of neuroscience 35 (2012): 203–225.

Calfa, Gaston, John J. Hablitz, and Lucas Pozzo-Miller. “Network hyperexcitability in hippocampal slices from Mecp2 mutant mice revealed by voltage-sensitive dye imaging.” Journal of neurophysiology 105.4 (2011): 1768.

Canolty, Ryan T., et al. “High gamma power is phase-locked to theta oscillations in human neocortex.” science 313.5793 (2006): 1626–1628.

Carlson, Greg C., and Douglas A. Coulter. “In vitro functional imaging in brain slices using fast voltage-sensitive dye imaging combined with whole-cell patch recording.” Nature protocols 3.2 (2008): 249–255.

Chahrour, Maria, and Huda Y. Zoghbi. “The story of Rett syndrome: from clinic to neurobiology.” Neuron 56.3 (2007): 422–437.

Chapleau, Christopher A., et al. “Hippocampal CA1 pyramidal neurons of Mecp2 mutant mice show a dendritic spine phenotype only in the presymptomatic stage.” Neural plasticity 2012 (2012).

Cohen, Lawrence B., and Brian M. Salzberg. Optical measurement of membrane potential. Springer Berlin Heidelberg, 1978.

Dani, Vardhan S., et al. “Reduced cortical activity due to a shift in the balance between excitation and inhibition in a mouse model of Rett syndrome.” Proceedings of the National Academy of Sciences of the United States of America 102.35 (2005): 12560–12565.

D'Cruz, Jennifer Anne, et al. “Alterations of cortical and hippocampal EEG activity in MeCP2-deficient mice.” Neurobiology of disease 38.1 (2010): 8–16.

De Filippis, B., L. Ricceri, and G. Laviola. “Early postnatal behavioral changes in the Mecp2-308 truncation mouse model of Rett syndrome.” Genes, Brain and Behavior 9.2 (2010): 213–223.

Ezeonwuka, Chinelo D., and Mojgan Rastegar. “MeCP2-Related Diseases and Animal Models.” Diseases 2.1 (2014): 45–70.

Gantz, Stephanie C., et al. “Loss of Mecp2 in substantia nigra dopamine neurons compromises the nigrostriatal pathway.” The Journal of Neuroscience 31.35 (2011): 12629–12637.

Gloveli, Tengis, et al. “Differential involvement of oriens/pyramidale interneurones in hippocampal network oscillations in vitro.” The Journal of physiology 562.1 (2005): 131–147.

Goffin, Darren, et al. “Rett syndrome mutation MeCP2 T158A disrupts DNA binding, protein stability and ERP responses.” Nature neuroscience 15.2 (2012): 274–283.

Guy, Jacky, et al. “A mouse Mecp2-null mutation causes neurological symptoms that mimic Rett syndrome.” Nature genetics 27.3 (2001): 322–326.

Isoda, K., et al. “Postnatal changes in serotonergic innervation to the hippocampus of methyl-CpG-binding protein 2-null mice.” Neuroscience 165.4 (2010): 1254–1260.

Kitchigina, Valentina F. “Theta oscillations and reactivity of hippocampal stratum oriens neurons.” The Scientific World Journal 10 (2010): 930–943.

Liao, Wenlin, et al. “MeCP2+/− mouse model of RTT reproduces auditory phenotypes associated with Rett syndrome and replicate select EEG endophenotypes of autism spectrum disorder.” Neurobiology of disease 46.1 (2012): 88–92.

Liao, Xiaogang, and Edgar T. Walters. “The use of elevated divalent cation solutions to isolate monosynaptic components of sensorimotor connections in Aplysia.” Journal of neuroscience methods 120.1 (2002): 45–54.

Maccaferri, Gianmaria. “Stratum oriens horizontal interneurone diversity and hippocampal network dynamics.” The Journal of physiology 562.1 (2005): 73–80.

McLeod, F., et al. “Reduced seizure threshold and altered network oscillatory properties in a mouse model of Rett syndrome.” Neuroscience 231 (2013): 195–205.

Medrihan, Lucian, et al. “Early defects of GABAergic synapses in the brain stem of a MeCP2 mouse model of Rett syndrome.” Journal of neurophysiology 99.1 (2008): 112.

Moretti, Paolo, et al. “Learning and memory and synaptic plasticity are impaired in a mouse model of Rett syndrome.” The Journal of neuroscience 26.1 (2006): 319–327.

Neul, Jeffrey L., and Huda Y. Zoghbi. “Rett syndrome: a prototypical neurodevelopmental disorder.” The Neuroscientist 10.2 (2004): 118–128.

Pelka, Gregory J., et al. “Mecp2 deficiency is associated with learning and cognitive deficits and altered gene activity in the hippocampal region of mice.” Brain 129.4 (2006): 887–898.

Ren, Jun, et al. “Anxiety-related mechanisms of respiratory dysfunction in a mouse model of Rett syndrome.” The Journal of Neuroscience 32.48 (2012): 17230–17240.

Reynaud, Alexandre, et al. “Linear model decomposition for voltage-sensitive dye imaging signals: Application in awake behaving monkey.” Neuroimage 54.2 (2011): 1196–1210.

Ricceri, Laura, Bianca De Filippis, and Giovanni Laviola. “Mouse models of Rett syndrome: from behavioural phenotyping to preclinical evaluation of new therapeutic approaches.” Behavioural pharmacology 19.5–6 (2008): 501–517.

Ricciardi, Sara, et al. “Reduced AKT/mTOR signaling and protein synthesis dysregulation in a Rett syndrome animal model.” Human molecular genetics20.6 (2011): 1182–1196.

Sivaramakrishnan, Shobhana, Jason Tait Sanchez, and Calum Alex Grimsley. “High concentrations of divalent cations isolate monosynaptic inputs from local circuits in the auditory midbrain.” Frontiers in neural circuits 7 (2013).

Shepherd, Gordon MG, and David M. Katz. “Synaptic microcircuit dysfunction in genetic models of neurodevelopmental disorders: focus on Mecp2 and Met.” Current opinion in neurobiology 21.6 (2011): 827–833.

Stearns, N. A., et al. “Behavioral and anatomical abnormalities in Mecp2 mutant mice: A model for Rett syndrome.” Neuroscience 146.3 (2007): 907–921.

Tetzchner, Stephen, et al. “Vision, cognition and developmental characteristics of girls and women with Rett syndrome.” Developmental Medicine & Child Neurology 38.3 (1996): 212–225.

Uhlhaas, Peter J., and Wolf Singer. “Abnormal neural oscillations and synchrony in schizophrenia.” Nature Reviews Neuroscience 11.2 (2010): 100–113.

Uhlhaas, Peter J., et al. “A new look at gamma? High-(> 60 Hz) γ-band activity in cortical networks: function, mechanisms and impairment.” Progress in biophysics and molecular biology 105.1 (2011): 14–28.

Wang, Xiao-Jing. “Neurophysiological and computational principles of cortical rhythms in cognition.” Physiological reviews 90.3 (2010): 1195.

Weng, S-M., et al. “Synaptic plasticity deficits in an experimental model of rett syndrome: long-term potentiation saturation and its pharmacological reversal.” Neuroscience 180 (2011): 314–321.

Whittington, Miles A., et al. “Multiple origins of the cortical gamma rhythm.” Developmental neurobiology 71.1 (2011): 92–106.

Wither, Robert G., et al. “Regional MeCP2 expression levels in the female MeCP2-deficient mouse brain correlate with specific behavioral impairments.” Experimental neurology 239 (2013): 49–59.

Yamanouchi, Hideo, Makiko Kaga, and Masataka Arima. “Abnormal cortical excitability in Rett syndrome.” Pediatric neurology 9.3 (1993): 202–206.

Zhang, Liang, et al. “The MeCP2-null mouse hippocampus displays altered basal inhibitory rhythms and is prone to hyperexcitability.” Hippocampus 18.3 (2008): 294–309.

